# Differential Expression of Immune Related Genes in Taste Buds of Fed and Fasted Mice

**DOI:** 10.1101/339911

**Authors:** SM Crosson, S Currlin, O Moskalenko, S Yegorova, CD Dotson, S Zolotukhin

## Abstract

To study the effects of feeding on taste receptor cell transcriptional regulation, we performed RNA-seq analysis of circumvallate taste buds isolated from mice before or after food consumption. Here we report and compare the taste bud transcriptomes obtained from food-deprived, satiated, and *ad libitum* fed control mice. Despite sample heterogeneity inherent to the whole taste bud transcriptome, bioinformatics analysis yielded 144 differentially expressed transcripts associated with immunity, cytoskeletal structure, and protein folding between these groups. We also profiled the transcriptome obtained from *ad libitum* fed control mice based on receptor related gene ontology terms, demonstrating a use of this dataset in the identification of novel TRC receptors. The data presented here suggest that transcriptional regulation of immune cytokine signaling occurs in TRCs shortly after meal consumption, though additional analysis is required to confirm this hypothesis.

## Introduction

Taste buds are comprised of specialized epithelial cells known as taste receptor cells (TRCs) (Chaudhari and Roper 2010). These TRCs can be classified into 3 categories referred to as Type I, II, and III based on their morphological characteristics (Murray 1993; Pumplin *et al.* 1997; Yee *et al.* 2001). Type I TRCs are the proposed sodium sensitive cells, detected via expression of epithelial sodium channels (ENaC), which also serve a glial-like supporting role in the bud (Bigiani 2001; Dvoryanchikov *et al.* 2009; Vandenbeuch *et al.* 2008). Type II TRCs express different G-protein coupled receptors (GPCRs), which transduce sweet, bitter, and umami taste qualities (Adler *et al.* 2000; Mueller *et al.* 2005; Nelson *et al.* 2002; Nelson *et al.* 2001; Zhang *et al.* 2003; Zhao *et al.* 2003). The final cell type, Type III TRCs, detect sour tastants via cytoplasmic changes in pH (Huang *et al.* 2008; Ishimaru *et al.* 2006; Tu *et al.* 2018). While many genetic and electrophysiological studies have been conducted on the gustatory system, substantially less data on the TRC transcriptome is available. With the exception of a recent study which used RNA-seq to profile the transcriptome of *Tas1r3* positive and Type III TRCs (Sukumaran *et al.* 2017), the few other TRC transcriptome studies used micro array technology as opposed to RNA-seq (Hevezi *et al.* 2009; Moyer *et al.* 2009). Acquisition of transcriptome data for all TRC subtypes is essential for the discovery of novel receptors, as well as obtaining a better understanding of TRC gene expression patterns during different circadian and digestive phases.

The functioning of the gustatory system is intimately linked with the functioning of the digestive system and as such, can influence and be influenced by the different phases of digestion. For example, during the cephalic phase of digestion, initial tastant detection by sweet responsive TRCs stimulates secretion of insulin from pancreatic islets (Tonosaki *et al.* 2007). Likewise, TRCs express receptors for a number of gastrointestinal peptides and hormones that are secreted into the circulation and saliva during the cephalic, gastric, and intestinal digestive phases (Awazawa *et al.* 2011; Elson *et al.* 2010; Hurtado *et al.* 2012; Kawai *et al.* 2000; La Sala *et al.* 2013; Martin *et al.* 2009; Martin *et al.* 2010; Shin *et al.* 2010). These include receptors for glucagon-like peptide 1 (GLP-1), peptide tyrosine-tyrosine (PYY), glucagon, and insulin among other modulatory peptides and hormones. Furthermore, satiety peptides PYY, GLP-1, and glucagon modulate the taste perception of specific taste qualities, presumably by interacting with cognate receptors expressed on TRCs (Elson *et al.* 2010; La Sala *et al.* 2013; Takai *et al.* 2015). While a modulatory effect of these peptides on taste perception has been demonstrated in animal models, these results are contextually dependent on the ingestive or sated state of the animals. In other words, levels of GI hormones known to influence the gustatory system are not static throughout the day. In the TRCs as well, there may be transcriptional or translational differences depending on the circadian, digestive, or ingestive phase of the animal. To gain a better understanding of the transcriptional differences in the peripheral gustatory system during periods of feeding or fasting, we performed RNA-seq on circumvallate taste buds isolated from wild-type (WT) C57BL6/J mice under different feeding regiments. Specifically, we performed RNA-seq on taste bud messenger RNA isolated from 3 groups of mice referred to a “satiated”, “hungry”, and *“ad lib”,* then identified differentially expressed genes between each group. We performed gene enrichment analysis of the differentially expressed genes and found enrichment in pathways associated with immunity, cytoskeletal structure, chemokines, chaperone proteins, and protease inhibitors. This suggests that these specific pathways may be regulated in TRCs in response to feeding, though further investigation into the role of these pathways is TRCs is needed.

## Materials and Methods

### Animals

WT C57BL6/J mice used in this study were bread in house, housed at 22-24 ^o^C with a 14/10 hr light/dark cycle, and given *ad libitum* access to food and water except where otherwise noted. In the 24 hrs prior to tissue collection for RNA-seq analysis, 2 groups of mice were placed on restricted feeding schedules; while a third control group, hear after referred to as “ad lib” (n=7), received *ad libitum* access to food throughout the experiment. The first group of food restricted mice, hear after referred to as “hungry” (n=7), were fasted for the 24 hrs prior to tissue collection. The second group of mice, hear after referred to as “satiated” (n=7), were fasted for 23 hrs, then given *ad libitum* access to food for 1 hr prior to tissue collection. All animals had *ad libitum* access to water throughout the experiments except where specifically noted. Only male mice, age 20-24 weeks, were used in order to eliminate influences of the estrus cycle on hormone levels and feeding behavior. As well, the average body weight of mice was kept relatively constant across each group. Procedures were done in accordance with principals set forth by the National Research Council’s guide for the Care and Use of Laboratory Animals. This study was approved by the University of Florida Institutional Animal Care and Use Committee (IACUC).

### Circumvallate Taste Bud Isolation

Circumvallate taste buds were isolated from mouse tongues in accordance with protocols reported by Huang et al. 2005 (Huang 2005, Huang 2016). Briefly, mice were euthanized by CO2 asphyxiation followed by cervical dislocation after which, tongues were dissected. Tongues were washed in Tyrode’s buffer (140 mM NaCl, 5 mM KCl, 2 mM CaCl_2_, 1 mM MgCl_2_ 10 mM HEPES, 10 mM glucose, 10 mM sodium pyruvate, and 5 mM NaHCO_3_ pH ~ 7.2) then, the lateral ends of the circumvallate papillae were each injected with 50 μl of protease cocktail (Collagenase A 1 mg/ml, Dispase II 2.5 mg/ml, Elastase 0.25 mg/ml, and DNAse I 0.5 mg/ml) and tongues were incubated in calcium/magnesium free Tyrode’s buffer. Tongues were pinned in Slygard gel, covered in Tyrode’s buffer, then using surgical scissors the circumvallate epithelium was removed, inverted, and pinned to a new Slygard gel. Circumvallate epithelia were incubated shortly with protease cocktail, washed with Tyrode’s buffer, then washed with calcium/magnesium free Tyrode’s buffer, before pinning in fresh Slygard gels and covering with Tyrode’s buffer containing 0.5 mg/ml DNAse I. Taste buds were collected by manual aspiration using 60-80 μm flame polished capillary pipettes, ejected onto Cell-Tak coated coverslips, then washed successively in droplets of Tyrode’s buffer. Taste buds from each animal were then collected into microfuge tubes for RNA extraction. All buds were collected 1 hr after onset of the dark cycle.

### RNA extraction and cDNA library generation

RNA was extracted from taste buds using the Absolutely RNA Nanoprep Kit (Stratagene) and treated with RNAseOUT (ThermoFisher) to preserve RNA integrity. Quality control and total RNA amounts were determined using a Bioanalyzer, and only samples with RNA integrity numbers (RIN) greater than 7 were used for cDNA synthesis and subsequent sequencing. Poly-adenylated RNA was converted to cDNA using SuperScript III Reverse Transcriptase (ThermoFisher) and oligo(dT) primers. A cDNA library for Illumina sequencing was then constructed using the RNA-seq Library Preparation Kit (Kappa Biosystems). The cDNA libraries were transferred to the University of Florida Interdisciplinary Center for Biotechnology Research (ICBR) for next generation sequencing.

### RNA-Seq data analysis

Single end RNA-seq was performed on an Illumina HiSeq 2000 with average read length of 75 bp. A total of 7 biological replicates (n=7) were used for each experimental group with an average of 30 million reads obtained for each animal. Quality control of fastq files was conducted using FastQC (Andrews) after which additional trimming of adapter sequences was done using Trimmomatic (Bolger *et al.* 2014). Trimmed reads were then mapped to the GRCm38 p.4 mouse reference genome, downloaded from Ensembl (Zerbino *et al.* 2018), using Tophat2 running the Bowtie2 short read alignment algorithm (Trapnell *et al.* 2010). Samtools was used to convert sequence alignment map (SAM) output files to binary alignment map (BAM) format and transcript assembly was completed using Cufflinks (Li *et al.* 2009; Trapnell *et al.* 2010). A combined gene transfer format (GTF) assembly file was generated for differential expression analysis, using Cuffmerge, containing the reference GTF genome as well as the Cufflinks output for each sample (Trapnell *et al.* 2010). Cuffdiff was used to determine pairwise differential gene expression between the various experimental groups, and the cummeRbund R package was used to generate graphical representations of the RNA-seq data set (Trapnell *et al.* 2012; Trapnell *et al.* 2010).

### DAVID Enrichment Analysis

We used DAVID Bioinformatics (DAVID) (Huang *et al.* 2009) to perform functional annotation enrichment analysis of the differentially expressed genes identified via Cuffdiff. DAVID identified annotation records for 128 of the 144 differentially expressed genes and used annotation information from the following databases for enrichment analysis: COG_ONTOLOGY, UP_KEYWORDS, UP_SEQ_FEATURE, GOTERM_BP_DIRECT, GOTERM_CC_DIRECT, GOTERM_MF_DIRECT, BIOCARTA, KEGG_PATHWAY, INTERPRO, PIR_SUPERFAMILY, and SMART. Using the functional annotation clustering program with high stringency classifications, similarity threshold of 0.85, multiple linkage threshold of 0.5, and an EASE enrichment threshold of 0.01, DAVID identified 5 enriched annotation clusters among the differentially expressed genes.

### Panther Gene Ontology Profiling

To profile the transcript expression of receptor related genes in wild-type circumvallate taste buds, we used Panther gene ontology (GO) (Mi *et al.* 2017; Mi *et al.* 2013; The Gene Ontology Consortium 2017). All transcripts expressed at a frequency greater or equal to 2 transcripts per million (TPM) were used for gene ontology profiling. Panther identified a total of 6022 genes which were then profiled by molecular function GO (3789 total) and protein class GO (3540 total) terms. Using cummeRbund and the ggplot2 package in R, we generated expression plots and GO pie charts depicting transcripts associated with the GO molecular function subclass: G-protein coupled receptors. The same graphs were generated for transcripts associated with the Protein class subcategories: G-protein coupled receptors, cytokine receptors, ligand-gated receptors, and protein kinase receptors.

## Results

### Biological Replicate Variability and TRC Validation

To assess the intergroup variation of gene expression, we used cummeRbund to determine the squared coefficient of variance (CV^2^) as well as the gene dispersion of each group. The CV^2^ for each group was relatively high, indicating a large degree of variance between biological replicates within groups (Figure 1, panel A). As well, each group was overdispersed, further indicating a high degree of variation between biological replicates (Figure 1, panel B). Both models of variance showed a higher degree of variability at lower transcript expression levels as expected (Yoon and Nam 2017). To confirm enrichment of TRC RNA in our dataset, we queried the expression levels of multiple TRC markers, corresponding to all TRC subtypes, and confirmed that markers for Type I, II, and III TRCs are indeed expressed (Figure 2). There were no significant differences in the expression levels of the queried TRC markers between groups.

**Figure 1.**
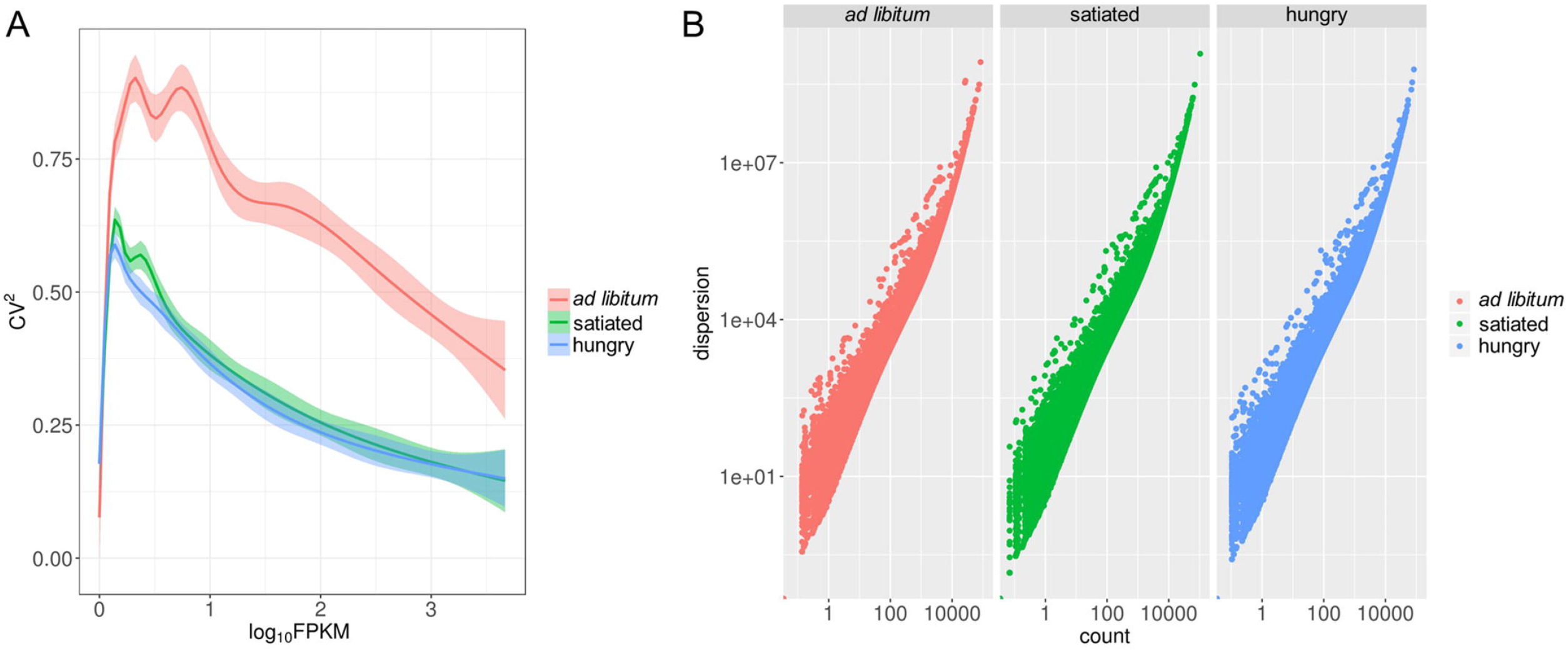
Gene expression variance within experimental groups. Both CV^2^ (A) and dispersion (B) models depict a high level of variance within each experimental group. Variance is higher for lower expressed transcripts as expected, and indicates heterogeneity between biological replicates.

**Figure 2.**
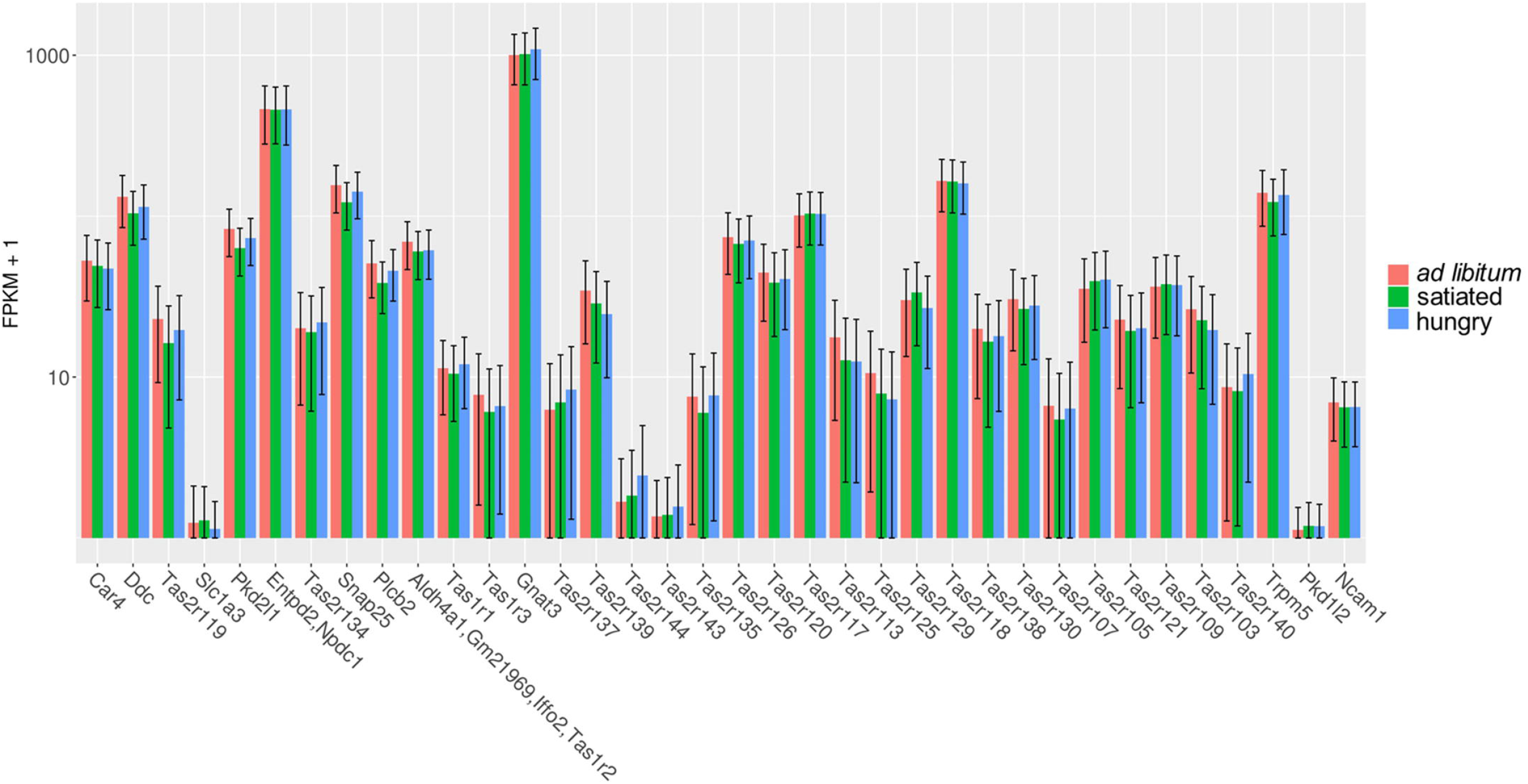
Type I, II, and III TRC markers are present in the RNA-seq dataset and expressed at similar levels in each experimental group. No significant differences were observed between groups for any of the TRC markers queried (q > 0.05).

### Differential Gene Expression between Ad Lib, Hungry, and Satiated Mice

Due to the high variance between biological replicates, we anticipated a relatively low number of differentially expressed genes between *ad lib,* hungry, and satiated mice. However, using a false discovery rate (FDR) of 5%, we identified a total of 144 differentially expressed genes between the 3 groups. Pairwise volcano plots indicate differentially expressed genes between *ad lib,* hungry, and satiated mice respectively (Figure 3). The significance expression matrix (Figure 4) displays the number of differentially expressed genes identified pairwise between each group. To look for patterns of gene expression across groups, genes were clustered by FPKM and expression was shown as a heatmap (Figure 5, panel A). Based on the heatmaps alone, there are only small clusters of genes with similar expression patterns. A heatmap showing fold change in FPKM of differentially expressed genes for hungry and satiated mice, relative to *ad lib* mice was constructed to observe changes in gene expression from the control state (Figure 5, panel B). There are small clusters of genes in satiated and hungry mice with opposing expression patterns, relative to *ad lib* mice, indicating that their expression may be associated with a pre or post-ingestive state (Figure 5). However, this type of comparison assumes that the *ad lib* mice are in an intermediary state relative to the satiated and hungry mice, which may not be the case.

**Figure 3.**
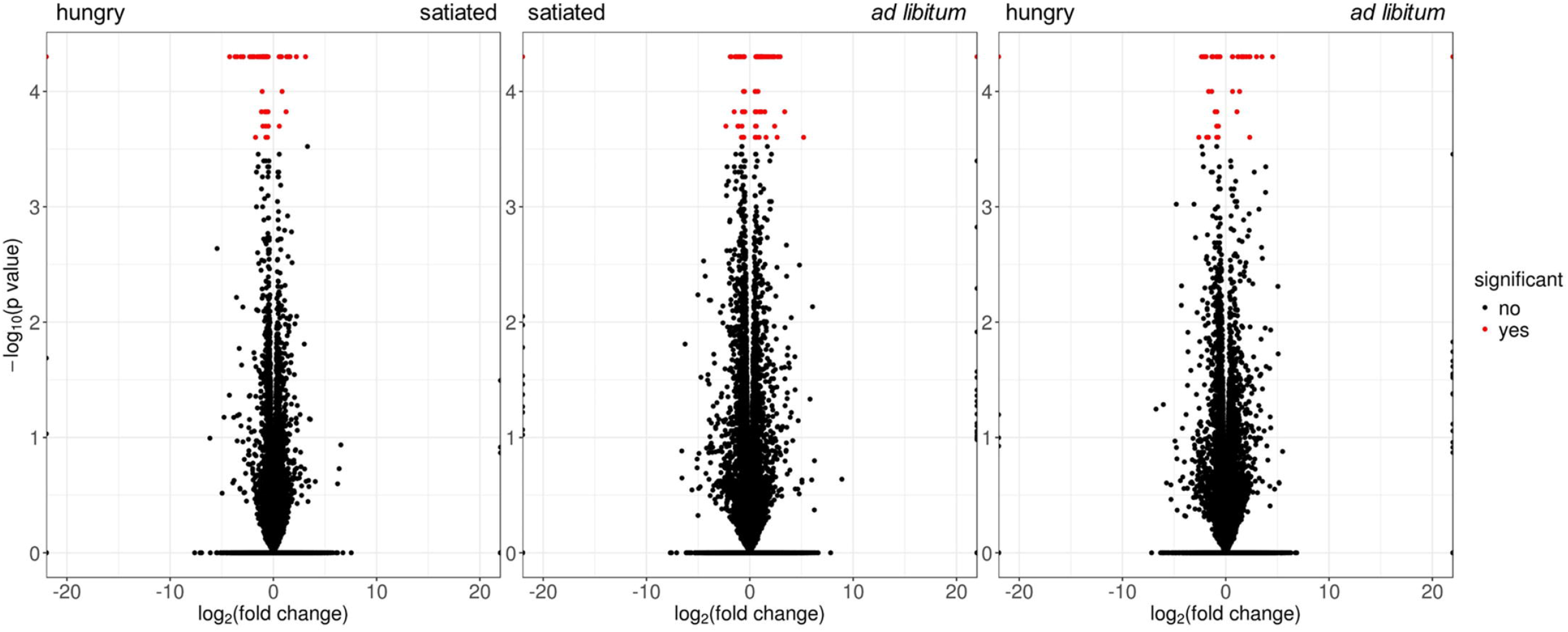
Volcano plots comparing gene expression levels between *ad lib,* hungry, and satiated mice. Genes that are significantly differentially expressed are shown in red (q < 0.05).

**Figure 4.**
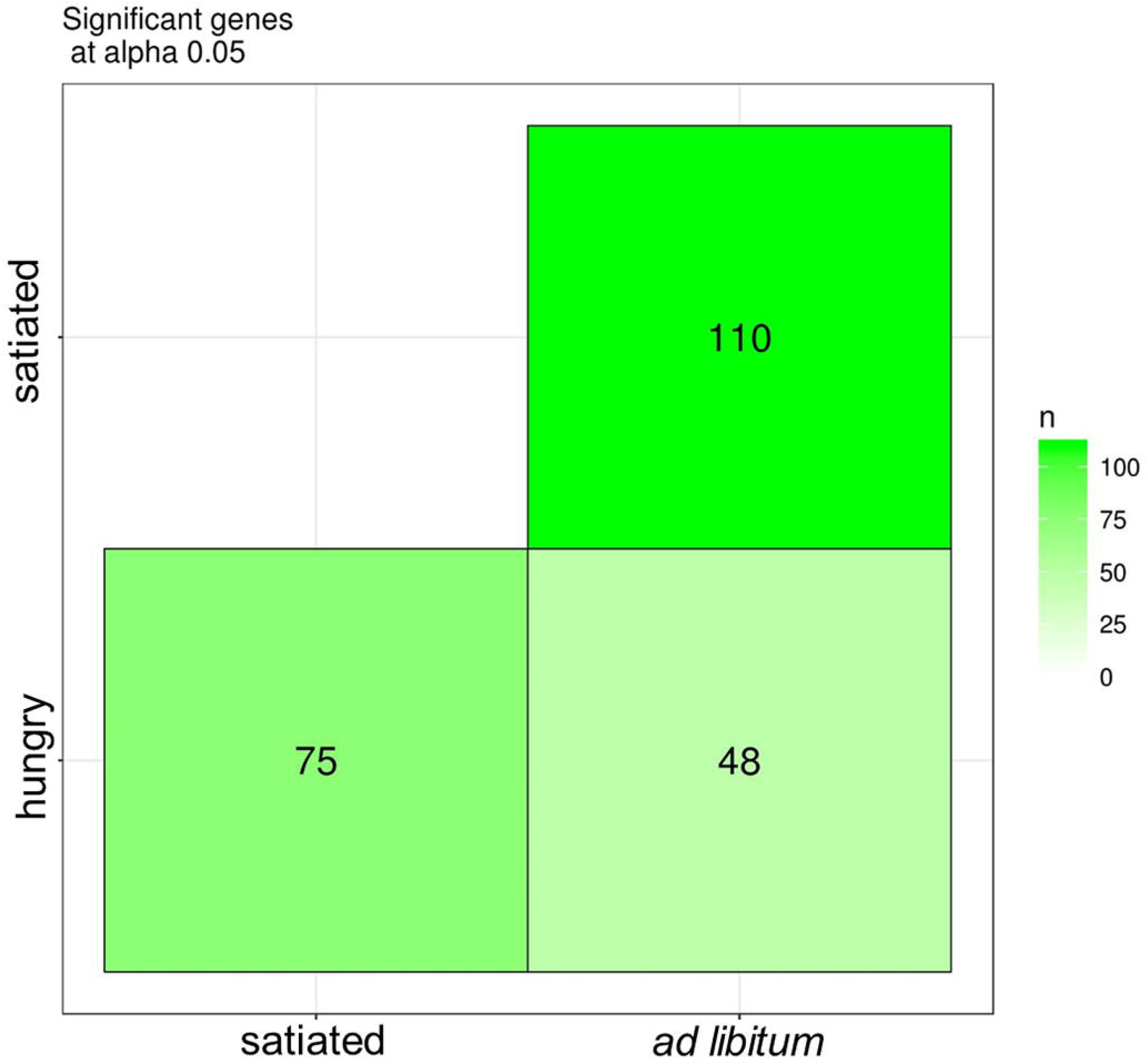
Differential expression matrix displaying the total number of differentially expressed genes between each experimental group (FDR 5%).

**Figure 5.**
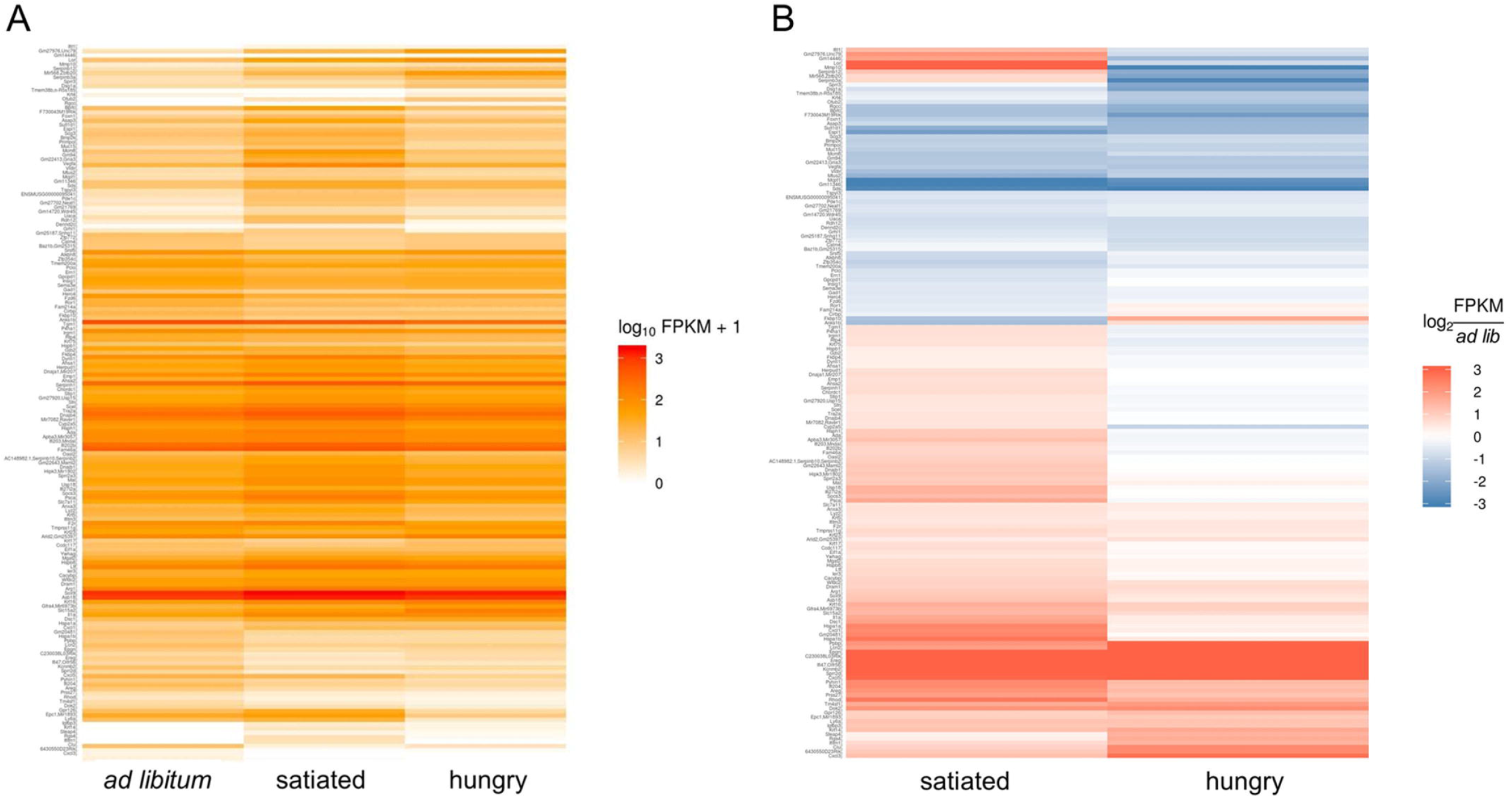
Gene expression heatmaps of the 144 differential genes. Heatmap of total gene expression (A) and fold change expression relative to the *ad lib* group (B), shows small clusters of genes with similar expression patterns.

### Functional Clustering of Differentially Expressed Genes using DAVID

To cluster differentially expressed genes based on their biological function, we used DAVID bioinformatics to perform functional annotation clustering using annotation information from the default DAVID databases (Huang *et al.* 2009). Using high stringency enrichment criteria with an EASE threshold of 0.01, DAVID identified 5 functional annotation clusters. Heatmaps depicting genes and associated annotation terms for each functional annotation cluster are shown in Figure 6. Functional annotation clusters 1-5 largely contain cytokeratin, chemokine, serpin protease inhibitor, immune related, and chaperone related genes, respectively (Figure 6). Expression levels of clustered genes were plotted across experimental groups to observe the pattern of gene expression changes (Figure 7). For each functional annotation cluster, most genes display the same expression pattern between *ad lib,* satiated, and hungry mice (Figure 7). Specifically, most enriched genes are upregulated in satiated mice relative to hungry or *ad lib* (Figure 7).

**Figure 6.**
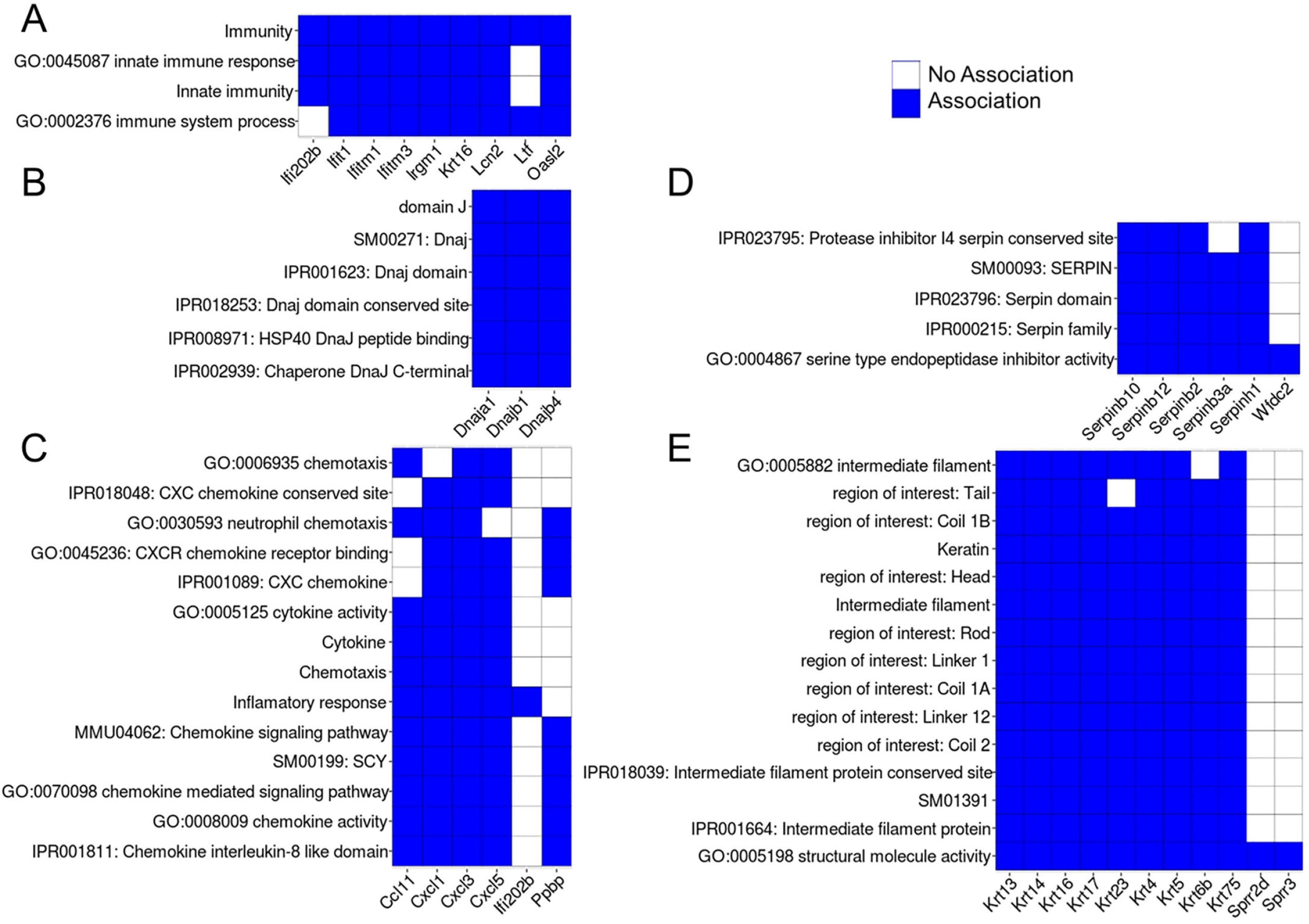
Functional annotation clustering of differentially expressed genes using DAVID bioinformatics. Heatmaps show association of genes in 5 different enrichment clusters with respective DAVID annotation terms. Blue cells denote an association with an annotation term, while white cells denote an absence of association.

**Figure 7.**
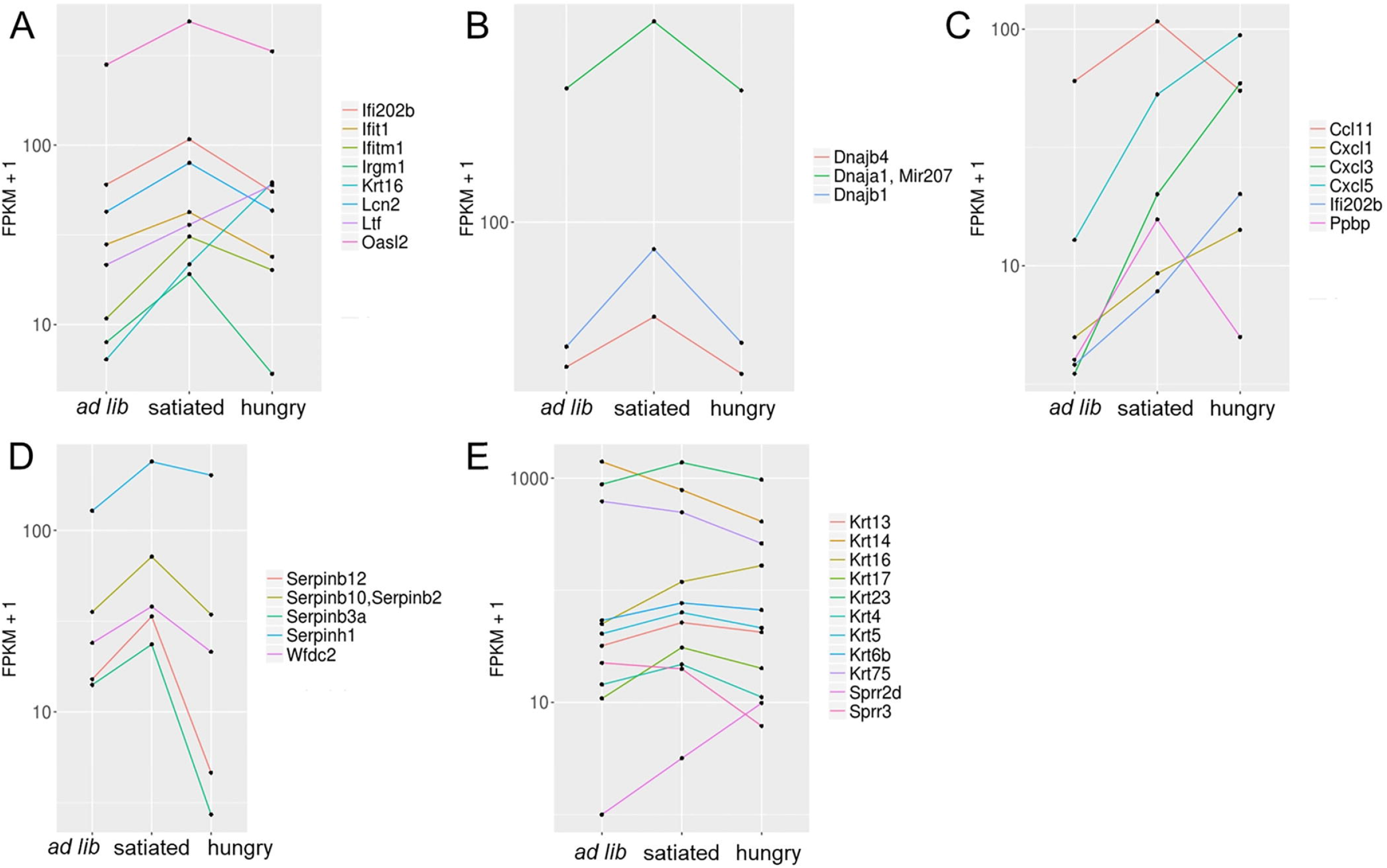
Gene expression levels of DAVID identified gene enrichment clusters for each group. Majority of genes are upregulated in satiated mice relative to *ad lib* and hungry mice except for genes in the chemokine enrichment cluster (C).

### Profiling of Taste Bud Transcriptome Using Keyword Gene Ontology Terms

Because of the limited transcriptome data available for murine taste buds, we thought it useful to determine an expression profile of transcripts associated with key GO annotations; specifically, those associated with receptor function. Using Panther gene ontology, we profiled the whole circumvallate taste bud transcriptome, from *ad lib* mice, for genes with expression levels greater than or equal to 2 transcripts per million (TPM). We specifically looked for the presence of genes with “receptor” associated GO terms. Pie charts, generated using Panther and the ggplot2 package in R, display either molecular function (3789 transcripts total, Figure 8 panel A) or protein class (3540 transcripts total, Figure 8 panel B) GO categories for the *ad lib* transcriptome at different annotation levels. Bar graphs depict expression of transcripts associated with receptor related GO terms, namely: GO:0004872, PC00084, PC00021, PC00141, and PC00194 (Figure 9). These data serve to validate previously reported findings, as well as identify uncharacterized receptors in the gustatory system which could serve as potential research targets.

**Figure 8.**
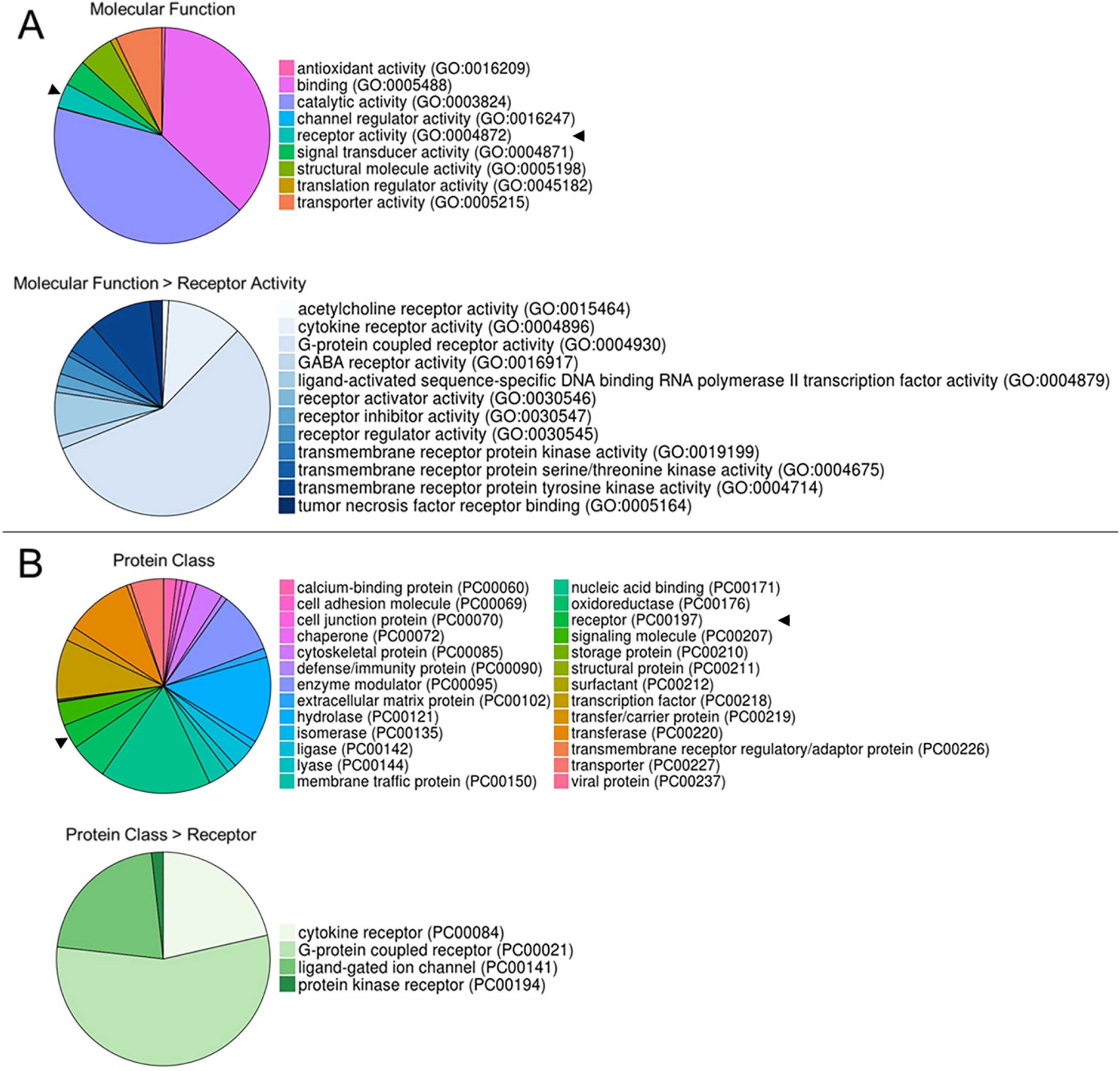
Panther gene ontology profiling of *ad lib* circumvallate taste bud transcriptome. Panel A displays Panther molecular function GO terms (3798 total transcripts), as well as functional GO terms associated with receptor activity (106 transcripts). Panel B contains Panther protein class GO terms (3540 total transcripts), as well as protein class GO terms associated with receptors (56 transcripts).

**Figure 9.**
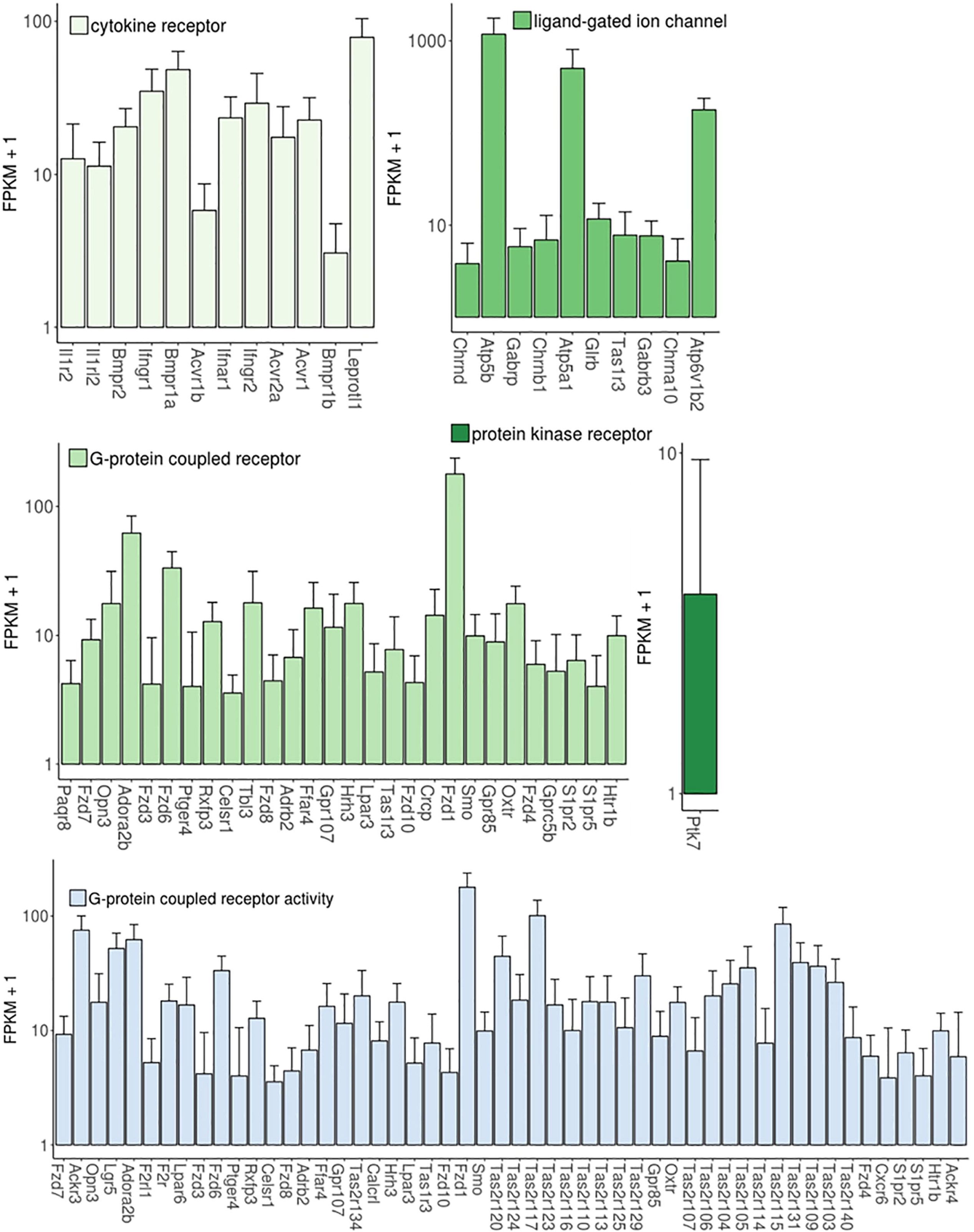
Expression levels of transcripts from *ad lib* circumvallate taste buds for select GO categories.

## Discussion

In this study, we used RNA-seq analysis to generate circumvallate taste bud transcriptomes from hungry and satiated mice. These transcriptomes contain RNA isolated from heterogeneous cell populations comprised primarily of Type I, II, and III TRCs. Because of the inherent heterogeneity present in the taste bud, as well as contamination with non-taste tissue present from taste bud isolation, we observed high sample variability as evident by dispersion and CV^2^ statistical analysis. Despite this variance, using a large number of biological replicates (n=7), we successfully identified 144 genes that are differentially expressed between satiated, hungry, and *ad lib* mice. Using gene enrichment analysis we identified 5 gene clusters associated with specific cellular pathways, which are differentially regulated between these groups. Additionally, we profiled the *ad lib* transcriptome based on receptor related GO terms in order to identify receptors novel to the gustatory system.

Gene enrichment analysis of differentially expressed genes indicated that pathways associated with chemokines, immunity, cytokeratins, serpin protease inhibitors, and chaperone proteins may be differentially regulated between hungry or sated animals. Differential regulation of chemokine and immune related transcripts between hungry and satiated mice is particularly intriguing, as immune cytokine signaling seems to be involved in taste cell communication (Feng *et al.* 2014b; Hevezi *et al.* 2009). Specifically, subpopulations of TRCs express interferon-γ, tumor necrosis factor, interlukin-10, and their cognate receptors (Feng *et al.* 2014a; Feng *et al.* 2012; Shi *et al.* 2012; Wang *et al.* 2007). They also express multiple types of toll-like receptors (Wang *et al.* 2009), which are responsible for detecting pathogenic microbial proteins. Furthermore, excessive inflammation in the taste bud is associated with taste dysfunction and increased TRC apoptosis, highlighting the importance of these pathways in taste bud homeostasis (Wang *et al.* 2009). Differential expression of cytokines and interferon inducible genes between hungry and satiated mice suggests that up or downregulation of these pathways, which are important for cell differentiation, proliferation, and survival (Feng *et al.* 2014b), may be induced shortly after meal consumption. This idea is further supported by the presence of cytokeratin proteins in the enrichment analysis, as some cytokeratins, such as K14 and K5, are expressed in subsets of taste progenitor cells (Okubo *et al.* 2009). Increased cell proliferation and differentiation may also explain the differential regulation of Hsp40 chaperone proteins, as these aid in protein folding and cellular transport (Cyr and Ramos 2015). However, further investigation into the role of these genes in TRC function is required to validate and support this hypothesis. Furthermore, we cannot exclude the possibility that some of the differentially expressed transcripts identified, may in fact be expressed in the contaminating non-taste tissue rather than in TRCs. It is likely that this would only be true for a small number of differentially expressed transcripts as the dataset is enriched for TRC RNA.

Because of the limited transcriptome data available for TRCs and the significance of receptors in TRC signaling, we used GO annotations to profile receptor expression in *ad lib* circumvallate taste buds. Various transcripts with G-protein coupled receptor functional annotations and/or receptor protein class annotations were present in the taste bud transcriptome. In addition to the traditional taste receptor genes, less studied receptors such as proposed fatty acid receptor *Ffar4* (Cartoni *et al.* 2010), as well as the orphan G-protein coupled receptor *GPR85* (Yuri *et al.* 2011), are also present. While mere detection of these receptors does not indicate functional significance, this serves to validate the presence of debated receptors such as Ffar4 and highlights receptors like the orphan *GPR85* which may play a novel role in the taste bud.

In summary we present here, circumvallate taste bud transcriptomes for fed, fasted, and *ad libitum* fed WT C57BL6 mice. Using gene enrichment analysis we showed that differentially expressed genes are enriched in pathways associated with immunity, chemokines, serine protease inhibitors, cytokeratin structural proteins, and chaperone proteins. We also profiled the *ad lib* transcriptome based on receptor related GO terms, in order to demonstrate the use of this transcriptome dataset in the identification of novel receptors. Overall, the data presented here suggest that specific cellular pathways may be regulated in TRCs with respect to food intake. However, further study is needed to understand the implications of these finding in a broader biological context.

## Acknowledgements

The authors would like to acknowledge the University of Florida’s Interdisciplinary Center for Biotechnology Research core facility for their assistance during the RNA-seq analysis.

## References

Adler E, Hoon MA, Mueller KL, Chandrashekar J, Ryba NJ, Zuker CS. 2000. A novel family of mammalian taste receptors. Cell 100: 693–702.

Andrews S. FastQC: a quality control tool for high throughput sequence data.

Awazawa M, Ueki K, Inabe K, Yamauchi T, Kubota N, Kaneko K, Kobayashi M, Iwane A, Sasako T, Okazaki Y, Ohsugi M, Takamoto I, Yamashita S, Asahara H, Akira S, Kasuga M, Kadowaki T. 2011. Adiponectin enhances insulin sensitivity by increasing hepatic IRS-2 expression via a macrophage-derived IL-6-dependent pathway. Cell Metab 13: 401–412.

Bigiani A. 2001. Mouse taste cells with glialike membrane properties. J Neurophysiol 85: 1552–1560.

Bolger AM, Lohse M, Usadel B. 2014. Trimmomatic: a flexible trimmer for Illumina sequence data. Bioinformatics 30: 2114–2120.

Cartoni C, Yasumatsu K, Ohkuri T, Shigemura N, Yoshida R, Godinot N, le Coutre J, Ninomiya Y, Damak S. 2010. Taste preference for fatty acids is mediated by GPR40 and GPR120. J Neurosci 30: 8376–8382.

Chaudhari N, Roper SD. 2010. The cell biology of taste. J Cell Biol 190: 285–296.

Cyr DM, Ramos CH. 2015. Specification of Hsp70 function by Type I and Type II Hsp40. Subcell Biochem 78: 91–102.

Dvoryanchikov G, Sinclair MS, Perea-Martinez I, Wang T, Chaudhari N. 2009. Inward rectifier channel, ROMK, is localized to the apical tips of glial-like cells in mouse taste buds. J Comp Neurol 517: 1–14.

Elson AE, Dotson CD, Egan JM, Munger SD. 2010. Glucagon signaling modulates sweet taste responsiveness. FASEB J 24: 3960–3969.

Feng P, Chai J, Zhou M, Simon N, Huang L, Wang H. 2014a. Interleukin-10 is produced by a specific subset of taste receptor cells and critical for maintaining structural integrity of mouse taste buds. J Neurosci 34: 2689–2701.

Feng P, Huang L, Wang H. 2014b. Taste bud homeostasis in health, disease, and aging. Chem Senses 39: 3–16.

Feng P, Zhao H, Chai J, Huang L, Wang H. 2012. Expression and secretion of TNF-α in mouse taste buds: a novel function of a specific subset of type II taste cells. PLoS One 7: e43140.

Hevezi P, Moyer BD, Lu M, Gao N, White E, Echeverri F, Kalabat D, Soto H, Laita B, Li C, Yeh SA, Zoller M, Zlotnik A. 2009. Genome-wide analysis of gene expression in primate taste buds reveals links to diverse processes. PLoS One 4: e6395.

Huang dW, Sherman BT, Lempicki RA. 2009. Systematic and integrative analysis of large gene lists using DAVID bioinformatics resources. Nat Protoc 4: 44–57.

Huang YA, Maruyama Y, Stimac R, Roper SD. 2008. Presynaptic (Type III) cells in mouse taste buds sense sour (acid) taste. J Physiol 586: 2903–2912.

Hurtado MD, Acosta A, Riveros PP, Baum BJ, Ukhanov K, Brown AR, Dotson CD, Herzog H, Zolotukhin S. 2012. Distribution of Y-receptors in murine lingual epithelia. PLoS One 7: e46358.

Ishimaru Y, Inada H, Kubota M, Zhuang H, Tominaga M, Matsunami H. 2006. Transient receptor potential family members PKD1L3 and PKD2L1 form a candidate sour taste receptor. Proc Natl Acad Sci U S A 103: 12569–12574.

Kawai K, Sugimoto K, Nakashima K, Miura H, Ninomiya Y. 2000. Leptin as a modulator of sweet taste sensitivities in mice. Proc Natl Acad Sci U S A 97: 11044–11049.

La Sala MS, Hurtado MD, Brown AR, Bohórquez DV, Liddle RA, Herzog H, Zolotukhin S, Dotson CD. 2013. Modulation of taste responsiveness by the satiation hormone peptide YY. FASEB J 27: 5022–5033.

Li H, Handsaker B, Wysoker A, Fennell T, Ruan J, Homer N, Marth G, Abecasis G, Durbin R, Subgroup GPDP. 2009. The Sequence Alignment/Map format and SAMtools. Bioinformatics 25: 2078–2079.

Martin B, Dotson CD, Shin YK, Ji S, Drucker DJ, Maudsley S, Munger SD. 2009. Modulation of taste sensitivity by GLP-1 signaling in taste buds. Ann N Y Acad Sci 1170: 98–101.

Martin B, Shin YK, White CM, Ji S, Kim W, Carlson OD, Napora JK, Chadwick W, Chapter M, Waschek JA, Mattson MP, Maudsley S, Egan JM. 2010. Vasoactive intestinal peptide-null mice demonstrate enhanced sweet taste preference, dysglycemia, and reduced taste bud leptin receptor expression. Diabetes 59: 1143–1152.

Mi H, Huang X, Muruganujan A, Tang H, Mills C, Kang D, Thomas PD. 2017. PANTHER version 11: expanded annotation data from Gene Ontology and Reactome pathways, and data analysis tool enhancements. Nucleic Acids Res 45: D183–D189.

Mi H, Muruganujan A, Casagrande JT, Thomas PD. 2013. Large-scale gene function analysis with the PANTHER classification system. Nat Protoc 8: 1551–1566.

Moyer BD, Hevezi P, Gao N, Lu M, Kalabat D, Soto H, Echeverri F, Laita B, Yeh SA, Zoller M, Zlotnik A. 2009. Expression of genes encoding multi-transmembrane proteins in specific primate taste cell populations. PLoS One 4: e7682.

Mueller KL, Hoon MA, Erlenbach I, Chandrashekar J, Zuker CS, Ryba NJ. 2005. The receptors and coding logic for bitter taste. Nature 434: 225–229.

Murray RG. 1993. Cellular relations in mouse circumvallate taste buds. Microsc Res Tech 26: 209–224.

Nelson G, Chandrashekar J, Hoon MA, Feng L, Zhao G, Ryba NJ, Zuker CS. 2002. An amino-acid taste receptor. Nature 416: 199–202.

Nelson G, Hoon MA, Chandrashekar J, Zhang Y, Ryba NJ, Zuker CS. 2001. Mammalian sweet taste receptors. Cell 106: 381–390.

Okubo T, Clark C, Hogan BL. 2009. Cell lineage mapping of taste bud cells and keratinocytes in the mouse tongue and soft palate. Stem Cells 27: 442–450.

Pumplin DW, Yu C, Smith DV. 1997. Light and dark cells of rat vallate taste buds are morphologically distinct cell types. J Comp Neurol 378: 389–410.

Shi L, He L, Sarvepalli P, McCluskey LP. 2012. Functional role for interleukin-1 in the injured peripheral taste system. J Neurosci Res 90: 816–830.

Shin YK, Martin B, Kim W, White CM, Ji S, Sun Y, Smith RG, Sévigny J, Tschöp MH, Maudsley S, Egan JM. 2010. Ghrelin is produced in taste cells and ghrelin receptor null mice show reduced taste responsivity to salty (NaCl) and sour (citric acid) tastants. PLoS One 5: e12729.

Sukumaran SK, Lewandowski BC, Qin Y, Kotha R, Bachmanov AA, Margolskee RF. 2017. Whole transcriptome profiling of taste bud cells. Sci Rep 7: 7595.

Takai S, Yasumatsu K, Inoue M, Iwata S, Yoshida R, Shigemura N, Yanagawa Y, Drucker DJ, Margolskee RF, Ninomiya Y. 2015. Glucagon-like peptide-1 is specifically involved in sweet taste transmission. FASEB J 29: 2268–2280.

The Gene Ontology Consortium. 2017. Expansion of the Gene Ontology knowledgebase and resources. Nucleic Acids Res 45: D331–D338.

Tonosaki K, Hori Y, Shimizu Y. 2007. Relationships between insulin release and taste. Biomed Res 28: 79–83.

Trapnell C, Roberts A, Goff L, Pertea G, Kim D, Kelley DR, Pimentel H, Salzberg SL, Rinn JL, Pachter L. 2012. Differential gene and transcript expression analysis of RNA-seq experiments with TopHat and Cufflinks. Nat Protoc 7: 562–578.

Trapnell C, Williams BA, Pertea G, Mortazavi A, Kwan G, van Baren MJ, Salzberg SL, Wold BJ, Pachter L. 2010. Transcript assembly and quantification by RNA-Seq reveals unannotated transcripts and isoform switching during cell differentiation. Nat Biotechnol 28: 511–515.

Tu YH, Cooper AJ, Teng B, Chang RB, Artiga DJ, Turner HN, Mulhall EM, Ye W, Smith AD, Liman ER. 2018. An evolutionarily conserved gene family encodes proton-selective ion channels. Science 359: 1047–1050.

Vandenbeuch A, Clapp TR, Kinnamon SC. 2008. Amiloride-sensitive channels in type I fungiform taste cells in mouse. BMC Neurosci 9: 1.

Wang H, Zhou M, Brand J, Huang L. 2007. Inflammation activates the interferon signaling pathways in taste bud cells. J Neurosci 27: 10703–10713.

Wang H, Zhou M, Brand J, Huang L. 2009. Inflammation and taste disorders: mechanisms in taste buds. Ann N Y Acad Sci 1170: 596–603.

Yee CL, Yang R, Böttger B, Finger TE, Kinnamon JC. 2001. “Type III” cells of rat taste buds: immunohistochemical and ultrastructural studies of neuron-specific enolase, protein gene product 9.5, and serotonin. J Comp Neurol 440: 97–108.

Yoon S, Nam D. 2017. Gene dispersion is the key determinant of the read count bias in differential expression analysis of RNA-seq data. BMC Genomics 18: 408.

Yuri M, Hiramoto M, Naito M, Matsumoto M, Matsumoto S, Morita S, Mori K, Yokota H, Teramura T. 2011. Identification and relative quantitation of an orphan G-protein coupled receptor SREB2 (GPR85) protein in tissue using a linear ion trap mass spectrometer. J Proteome Res 10: 2658–2663.

Zerbino DR, Achuthan P, Akanni W, Amode MR, Barrell D, Bhai J, Billis K, Cummins C, Gall A, Girón CG, Gil L, Gordon L, Haggerty L, Haskell E, Hourlier T, Izuogu OG, Janacek SH, Juettemann T, To JK, Laird MR, Lavidas I, Liu Z, Loveland JE, Maurel T, McLaren W, Moore B, Mudge J, Murphy DN, Newman V, Nuhn M, Ogeh D, Ong CK, Parker A, Patricio M, Riat HS, Schuilenburg H, Sheppard D, Sparrow H, Taylor K, Thormann A, Vullo A, Walts B, Zadissa A, Frankish A, Hunt SE, Kostadima M, Langridge N, Martin FJ, Muffato M, Perry E, Ruffier M, Staines DM, Trevanion SJ, Aken BL, Cunningham F, Yates A, Flicek P. 2018. Ensembl 2018. Nucleic Acids Res 46: D754–D761.

Zhang Y, Hoon MA, Chandrashekar J, Mueller KL, Cook B, Wu D, Zuker CS, Ryba NJ. 2003. Coding of sweet, bitter, and umami tastes: different receptor cells sharing similar signaling pathways. Cell 112: 293–301.

Zhao GQ, Zhang Y, Hoon MA, Chandrashekar J, Erlenbach I, Ryba NJ, Zuker CS. 2003. The receptors for mammalian sweet and umami taste. Cell 115: 255–266.

